# Production of diverse retinal analogues in engineered *Escherichia coli* through promiscuous carotenoid cleavage by Blh

**DOI:** 10.64898/2026.08.06.743266

**Authors:** Maiko Furubayashi

## Abstract

Nature produces hundreds of carotenoids, yet only a handful of the apocarotenoids derived from them are accessible through microbial production. The best-known example is retinal, the chromophore of rhodopsins and a precursor of pharmaceutical retinoids, which is generated by the central cleavage of β-carotene. Whether the same cleavage chemistry can be extended to other carotenoids, yielding retinal analogues that differ in their ring structures, and potentially in their biological activities, has remained largely untested. In this study, we demonstrate a pathway engineering approach in *E. coli* for the biosynthesis of diverse retinal analogues by leveraging substrate promiscuity of Blh, a bacterial carotenoid cleavage enzyme originally identified in microbial rhodopsin gene clusters. While initial co-expression of Blh with carotenoid pathway genes often resulted in the production of retinal (by cleavage of β-carotene intermediate), we found that by optimizing the expression level of Blh, carotenoids such as astaxanthin or canthaxanthin were cleaved efficiently. Structure-guided engineering of Blh, informed by its predicted substrate-binding cavity, further improved the cleavage of zeaxanthin. This expanded catalytic activity suggests that Blh can serve as a versatile biocatalyst for the production of diverse retinal analogues, potentially yielding compounds with a range of biological activities. Furthermore, our findings raise the possibility of diverse biological roles for these enzymes in their native biological contexts.

**Importance:** This study demonstrated the successful biosynthesis of a diverse array of retinal analogues in engineered *Escherichia coli* through the heterologous expression of Blh, a β-carotene cleavage dioxygenase, together with several carotenoid pathways. Careful design of the Blh expression construct enabled modulation of retinoid proportions in the engineered pathway. This work uncovers previously unrecognized substrate promiscuity of Blh, revealing its capacity to accept carotenoids beyond β-carotene as substrates. For the first time, the predicted structure of Blh revealed the enzyme’s substrate cavity. Rational engineering by amino acid substitution designed to expand the cavity enabled the improved cleavage of hydroxylated carotenoids. These findings open new avenues for both fundamental research and biotechnological applications and have the potential to impact the microbial production of valuable retinoids.

## Introduction

Retinal (**Fig. 1**), the aldehyde form of vitamin A, is a crucial molecule that functions as a chromophore in both animal and microbial rhodopsins^1,2^, and is also a key precursor to a diverse group of molecules known as retinoids. Naturally occurring retinoids play crucial roles in regulating fundamental biological processes, including development, reproduction, immune function, and vision,^3^ while many synthetic retinoids are widely used as pharmaceutical compounds for skin health or cancer treatment.^3–5^ The diverse roles of retinoids highlight their biotechnological potential, driving increasing interest in their microbial production.^6–12^

**Figure 1.**
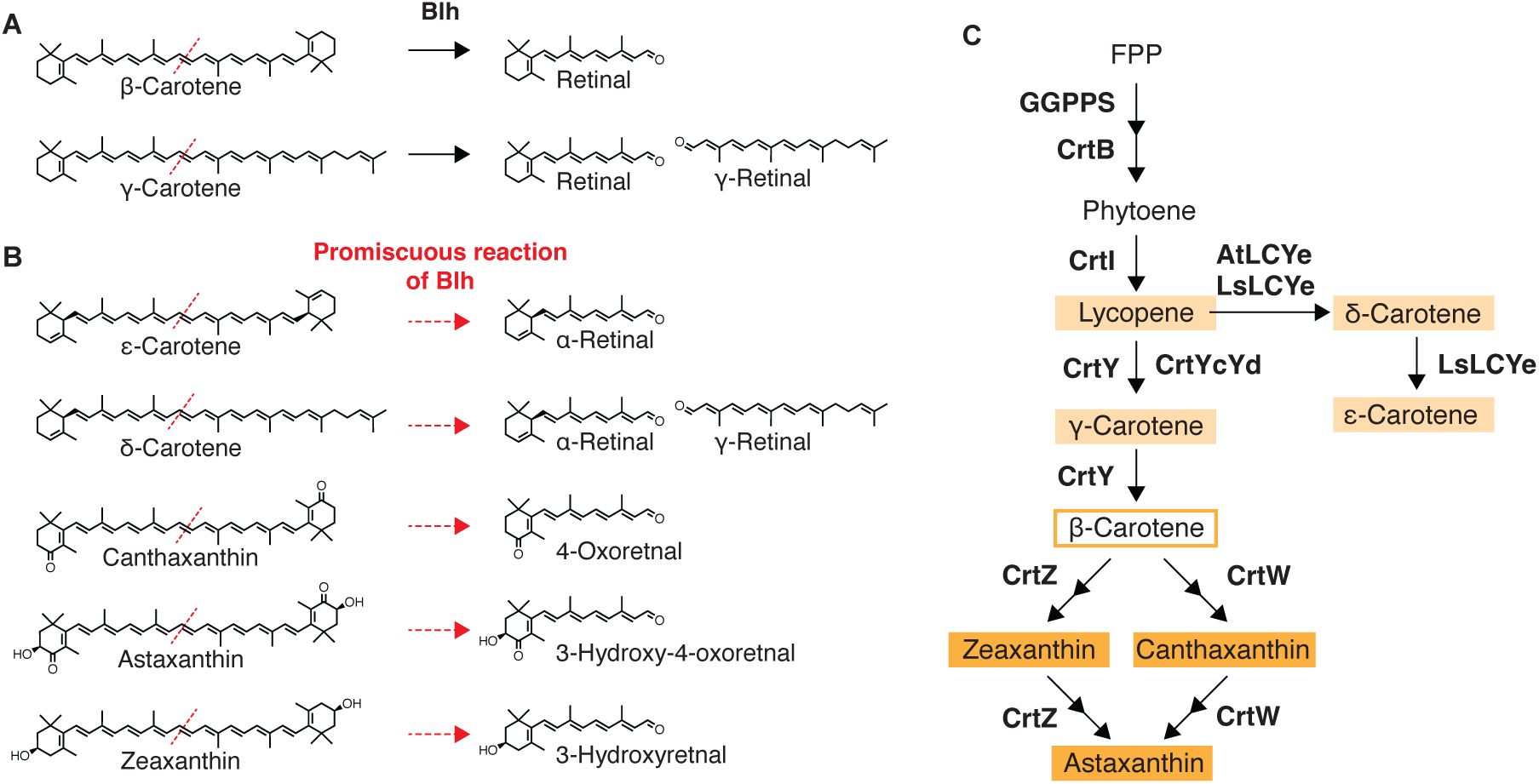
Blh-mediated production of retinoids in this study. (**A**) Previously known wildtype Blh activity. (**B**) Promiscuous Blh activity identified in this study. (**C**) Reconstructed carotenoid/retinoid pathway in *E. coli*. Enzymes are shown in bold face. Carotenoids upstream β-carotene are indicated in light orange, while the β-carotene derived carotenoids are indicated in dark orange.

Retinal is biosynthesized through the central (15,15’) cleavage of β-carotene, representing one of the most important “apocarotenoids” –a diverse class of molecules derived from carotenoid cleavage.^13–15^ Carotenoid cleavage enzymes catalyze the biosynthesis of apocarotenoids including retinal. Two structurally unrelated families of these enzymes exist in nature, and the term “carotenoid cleavage dioxygenases” (CCD) is conventionally applied only to the first. CCDs constitute a superfamily^16^ of protein homologous to the animal β-carotene 15,15’-dioxygenase 1 (BCO1)^17,18^, and to many plant enzymes (e.g. CCD1, CCD4, CCD7/8, NCED);^15^ they are soluble proteins with a 7-bladed β-propeller fold. BCO1-type CCDs are widely distributed and conserved across animals and plants,^16,19^ and have also been identified in some bacteria or archaea^20,21^. This family shows remarkable diversity in both substrate/product specificity and cleavage position: while BCO1 itself cleaves β-carotene at the 15,15’-position to synthesize retinal, its homologs such as BCO2^22^ or plant CCDs can cleave various carotenoids at different positions.

The second family, which is the focus of this study, cleaves carotenoids by the same dioxygenase mechanism,^23^ but shares no sequence or structural similarity with the CCD superfamily. The enzymes are named Brp (“Bacterioopsin-Related Protein”^24^) and Blh (“Brp-like homolog”^25^), which were first discovered in association with microbial rhodopsin gene clusters.^26,27^ While this enzyme family had long been only reported from bacteria and archaea, in relation to the microbial rhodopsin gene clusters,^28–33^ a recent genomic analysis^28^ has revealed the presence of Brp/Blh homologs and domains in eukaryotic lineages (such as protists, phototrophic eukaryotes, fungi, arthropods, and amoebozoans) via transposon-mediated transfer from bacteria/archaea. Unlike the BCO1-type CCDs that have been extensively biochemically characterized, the Blh/Brp-family enzymes only have a single comprehensive *in vitro* study^23^ for substrate specificity, suggesting activity primarily toward β-carotene and related compounds containing β-rings. It has been known that many bacteria harboring *brp*/*blh* genes also produce diverse carotenoids beyond β-carotene, the substrate for retinal, raising questions about substrate specificities for Blh. Since the interest in microbial production of retinoid and its derivatives is increasing, Brp/Blh can be a good candidate with much undiscovered potential.

In this study, we aimed to establish a platform for the production of diverse retinal analogue compounds by co-expressing bacterial Blh enzymes with various carotenoid biosynthetic pathways in an engineered *Escherichia coli* (*E. coli*) system. This approach allowed us to generate various retinal analogues, demonstrating that Blh can cleave not only β-carotene but also several other carotenoids, whose cleavage has not been reported elsewhere, such as canthaxanthin, astaxanthin, and zeaxanthin. The platform together with the AlphaFold2 prediction of protein structure led us to investigate the enzyme substrate cavity, enabling us to find a mutation that improves zeaxanthin (3-hydroxy-β-end) cleavage. These findings reveal unexpected substrate promiscuity of Blh and highlight its potential for biotechnological applications. Our findings may also suggest possible new roles for these enzymes in their native biological contexts.

## Results

### 1. Construction of Plasmids for β-Carotene Cleavage by Blh in *E. coli*

Among the Blh enzymes that have been used for retinal biosynthesis in *E. coli*^6–8,34–36^, for this study, Blh from uncultured marine bacterium 66A03^23^ was selected, as it demonstrated superior retinal production^6^ and was the only variant characterized for substrate specificity *in vitro*^23^.

To establish a retinal production system using the Blh enzyme in *E. coli*, two-plasmid system was designed, with one plasmid encoding the *blh* gene and the other encoding the carotenoid operons. Previous studies^6,7^ demonstrated that retinal production in *E. coli* was achievable by a similar co-expression strategy; however, these studies also revealed that a portion of the produced retinal was subsequently converted to retinol (an alcohol form of retinal) by *E. coli* endogenous enzymes, and when vectors encoding a chloramphenicol (Cm) resistance gene (Cm acetyltransferase) were used, retinal underwent acetylation to form retinyl acetate. To minimize the acetylated byproduct formation in this study, both Cm and kanamycin (Km) vectors were initially constructed as follows.

The carotenoid operon was cloned into p15A-based Cm or Km vectors (designated as pACm or pAKm, respectively) (**Supplementary Table 1** and **Supplementary Fig. 1 and 2**). The *blh* gene was positioned under the control of the P*_tac_* promoter in pUC ori vectors with an ampicillin resistance marker. The ribosome binding sites (RBSs) for Blh were designed and calculated using an RBS calculator^37^ from De Novo DNA (www.denovodna.com) (See **Supplementary Table 1** for the RBS sequence and the corresponding translation initiation rate [TIR].) The in-house strong RBS (calculated TIR of 15549) showed impaired growth suggesting the expression level was too high, so the medium RBS (TIR aimed for 2000, actual 1878) and weak RBS (TIR aimed for 100, actual 135) were designed and named as pUC-P_tac_-R_m_-Blh and pUC-P_tac_-R_w_-Blh, respectively. The β-carotene plasmids (pAKm-Beta or pACm-Beta) and pUC-P_tac_-R_w_-Blh were co-introduced into *E. coli* and cultured in a 10-mL media using 50-mL tubes. Dodecane was added (10% (v/v)) to the culture medium as an upper layer to facilitate retinal partitioning during cultivation.^6^

**Figure 2.**
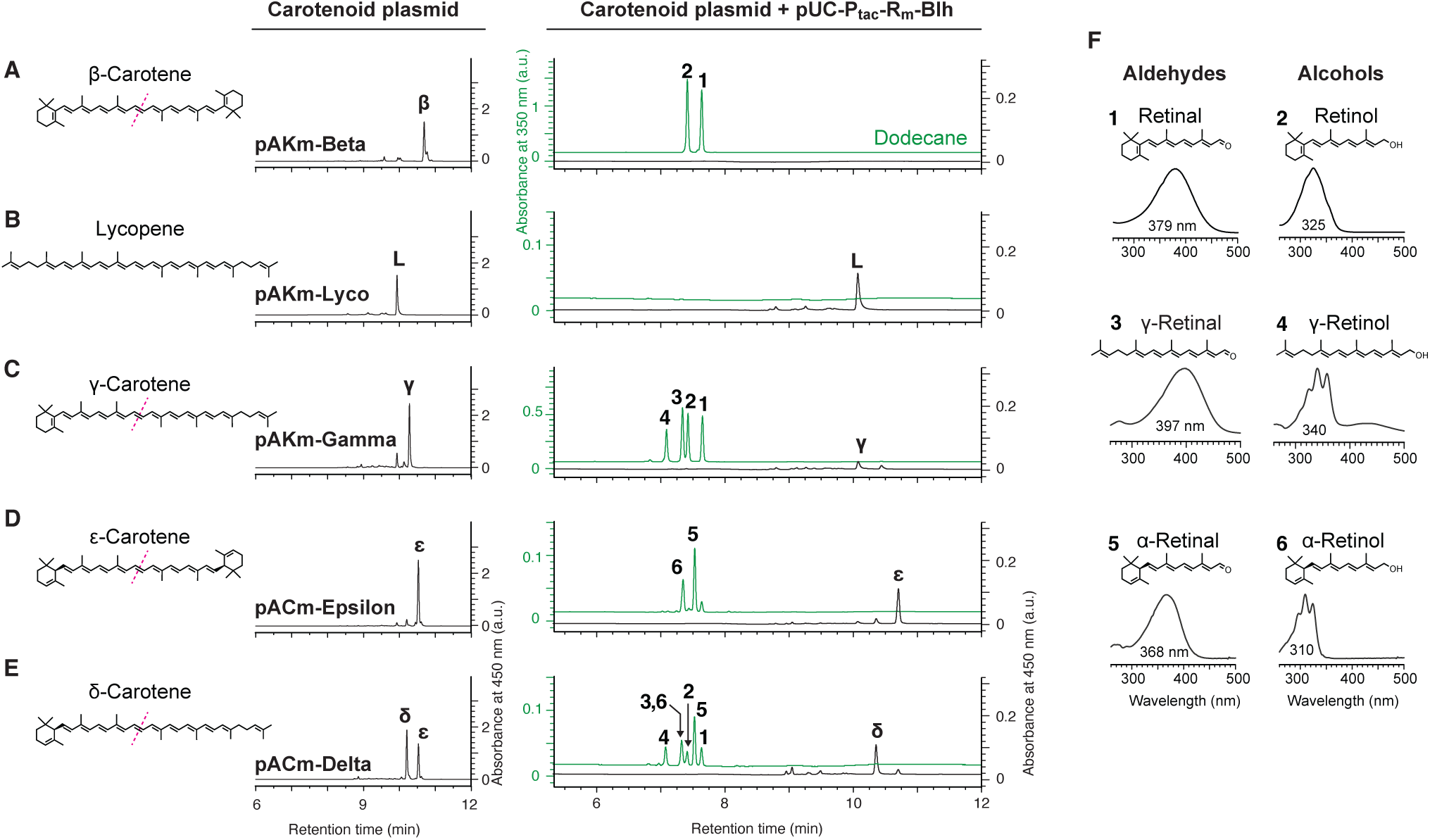
Blh-catalyzed production of diverse retinoids from non-β-carotene derived carotenoids (and a β-carotene control) in *E. coli*. Retinoid and carotenoid analysis was performed on extracts from *E. coli* cultures harboring various carotenoid biosynthesis plasmids (see **Supplementary** Fig. 1 **and Supplementary Table 1**), with or without Blh expression. For each panel, the chemical structure of the target carotenoid is shown on the left. Adjacent to it are two chromatograms of *E. coli* extracts without (left) or with (right) pUC-P_tac_-R_m_-Blh plasmid. In the chromatogram charts, the carotenoid fraction (acetone extract from pellet) is shown in black, while the retinoid fraction (dodecane overlay) is shown in green. The carotenoid biosynthesis plasmids used, and their corresponding carotenoids, are as follows: (**A**) pAKm-Beta for β-carotene, (**B**) pACm-Lyco for lycopene, (**C**) pAKm-Gamma for γ-carotene, (**D**) pACm-Epsilon for ε-carotene, (**E**) pACm-Delta for δ-carotene. (**F**) The absorbance spectrum and corresponding chemical structure of the compound of each major peak.

Expression of Blh in *E. coli* harboring pAKm-Beta (Km^R^ marker) resulted in the production of retinal and retinol in the upper dodecane layer (**Fig. 2A** and **Supplementary Fig. 2C**), with minimal residual β-carotene detected in the cell pellets. In contrast, Blh expression in *E. coli* containing pACm-Beta (Cm^R^ marker), yielded retinal and retinol, along with a minor peak corresponding to retinyl acetate (**Supplementary Fig. 2E**). Previous reports by Jang et al.^6,7^ documented retinyl acetate accounted for 10-20% (in mg/L) of the total product, whereas our results showed significantly lower levels of 3.5%. This disparity may be attributed to their enhanced upstream pathway for increased carotenoid/retinoids production.^6,7^ Given the negligible retinyl acetate formation observed in our study, both Cm^R^ and Km^R^ vectors were deemed suitable for subsequent experiments. The proportion of retinol produced was approximately 40% of retinal for both vectors, likely resulting from endogenous enzyme activity.^6,7^

### 2. Production of retinoids from non-β-carotene-derived carotenoids

Blh has been used in microbial production of retinal/retinol by co-expression with the β-carotene pathway; however, its substrate range beyond β-carotene cleavage remains largely unexplored, with no studies to date having tested its co-expression with other carotenoid pathways. The only comprehensive study examining the substrate specificity of Blh was an *in vitro* analysis by Kim et al.^23^ Their work, using a purified, soluble form of Blh (despite its prediction as a membrane protein), demonstrated that while Blh could cleave carotenoids containing a β-ring on at least one side of the chemical structure (γ-carotene, α-carotene, β-cryptoxanthin and β-apo-4’-carotenal), it did not cleave other carotenoids such as lycopene, zeaxanthin, lutein, and shorter apocarotenoids (such as β-apo-8’-carotenal and β-apo-12’-carotenal). However, these *in vitro* results were obtained using a solubilized form of the enzyme with detergent-solubilized substrates, conditions that may differ considerably from the cellular membrane environment. Here, we co-expressed Blh with our previously constructed carotenoid biosynthetic pathways^38^ in *E. coli* to produce retinoids from non-β-carotene carotenoids.

Initially, we focused on carotenoid pathways that do not involve β-carotene as an intermediate, specifically those yielding lycopene, γ-carotene, δ-carotene, ε-carotene (**Fig. 1C** and **Fig. 2**). First, introduction of P_tac_-R_m_-Blh into lycopene-producing strains showed negligible cleavage product formation (**Fig. 2B**), in agreement with the previous *in vitro* study.^23^ Next, expression of Blh in γ-carotene-producing *E. coli* resulted in the detection of retinal, retinol, γ-retinal and γ-retinol (**3** and **4** in **Fig. 2C**) in the dodecane layer, indicating that γ-carotene could be cleaved by Blh. This observation, too, corroborated previous *in vitro* findings^23^. Subsequently, expression of Blh in ε-carotene-producing *E. coli* resulted in two new peaks (**5** and **6** in **Fig. 2D**), alongside residual ε-carotene in the cell pellet. The absorbance spectrum of **5** and **6** showed a blue-shifted absorbance maximum relative to those of retinal and retinol, which indicated α-retinal and its alcohol derivative α-retinol, since the double bond in the ε-ring is isolated, resulting in a shorter conjugation length. Finally, in δ-carotene-producing strains (containing an ε-ring on one terminus and a linear ψ-end structure on the other), Blh co-expression led to the detection of α-retinal, γ-retinal, and their alcohol derivatives (**Fig. 2E**).

These results represent the first demonstration of γ-retinal/γ-retinol and α-retinal/α-retinol production in *E. coli*. Notably, the successful production of α-retinal/α-retinol from ε-carotene was unexpected, as the previous in vitro study^23^ had only shown cleavage of substrates containing a β-ring moiety. Our results indicate that Blh can also accept the ε-ring moiety as a substrate (Fig. 1B, top two diagrams), albeit with reduced efficiency compared to β-ring substrates, further expanding the range of retinoids accessible through microbial production.

### 3. Production of retinoids from β-carotene-derived carotenoids

We further explored the Blh-mediated production of retinoids from the β-carotene-derived carotenoids, specifically zeaxanthin, canthaxanthin and astaxanthin (**Fig. 1C** and **Fig. 3**). Notably, the cleavage of canthaxanthin and astaxanthin by Blh has not been reported to date. For all samples, carotenoid production (from samples without Blh coexpression) resulted in a single carotenoid product (**Fig. 3A-D**, left chromatogram). Blh co-expression by introducing pUC-P_tac_-R_m_-Blh to these strains resulted in consuming all carotenoids to produce retinoids (**Fig. 3A-D**, right chromatogram). Here, while the retinal (**1**) and retinol (**2**) (β-carotene cleavage products) were the primary products, several novel peaks (**7, 8, 9**) were observed in the chromatogram charts. Based on the absorbance spectrum (lambda max at 381 nm), mass spectra and product ion spectra (**Fig. 3G-I**), these peaks could be identified as 15,15’-cleavage of zeaxanthin, canthaxanthin and astaxanthin, i.e. 3-hydroxy-retinal (**7**) (hereafter referred to as “zea-retinal”), 4-oxo-retinal (**8**) (“cantha-retinal”) and 3-hydroxy-4-oxo-retinal (**9**) (“asta-retinal”), respectively. For peaks **8** and **9**, we observed additional slightly smaller peaks after 0.2 min, which might be cis isomers, since they showed a close absorbance maximum (380 nm) and same mass spectrum (indicated as **8’** and **9’**).

**Figure 3.**
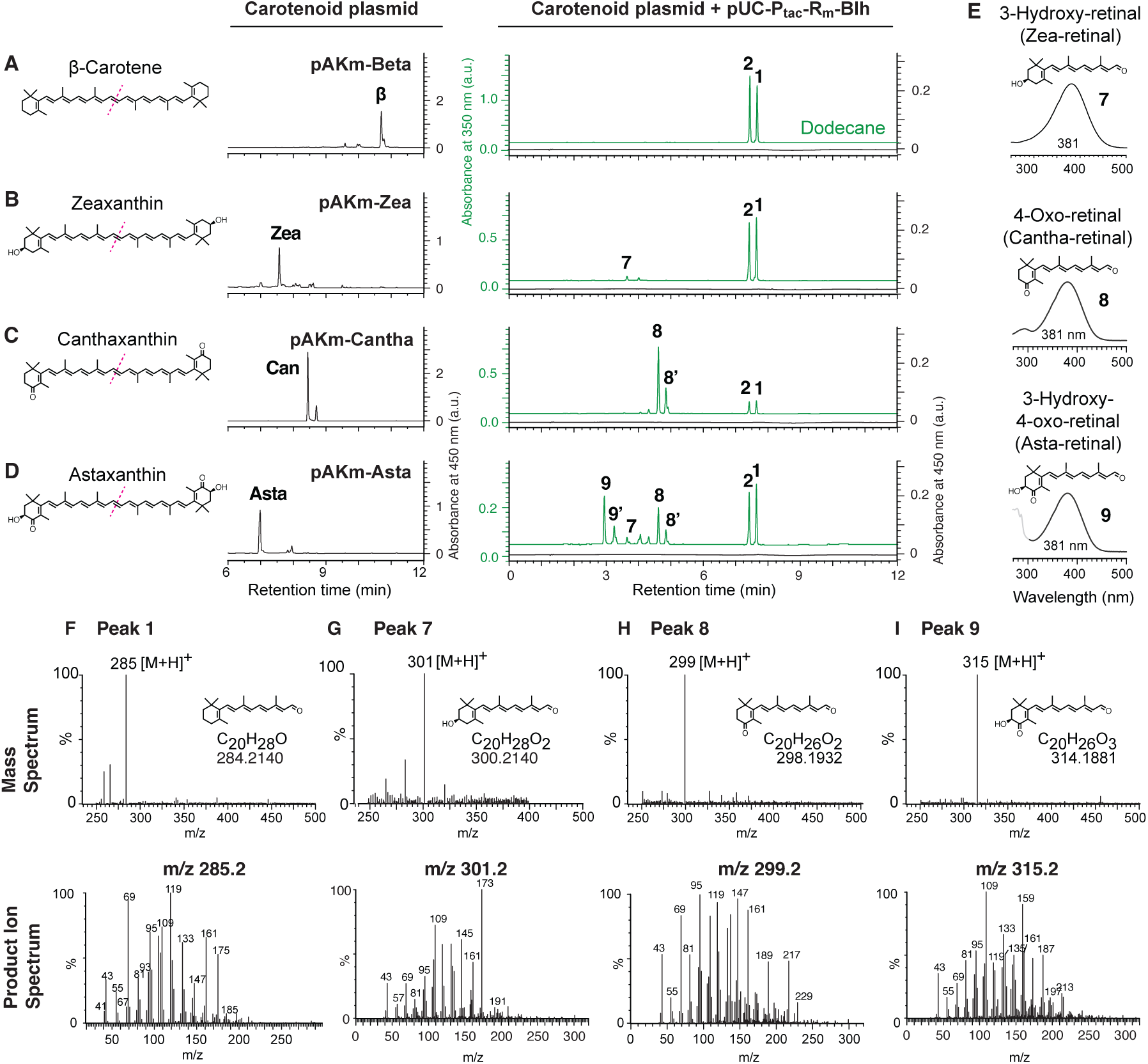
Blh-catalyzed production of diverse retinoids from β-carotene derived carotenoids in *E. coli*. Retinoid and carotenoid analysis was performed on extracts from *E. coli* cultures harboring various carotenoid biosynthesis plasmids, with or without Blh expression. Similar to Figure 2, panels (**A-D**) each show the chemical structure of the target carotenoid, two chromatograms (without and with Blh expression). In these chromatogram charts, the carotenoid fraction is shown in black, while the retinoid fraction (dodecane overlay) in green. The carotenoid biosynthesis plasmids used, and their corresponding carotenoids, are as follows: (**A**) pAKm-Beta for β-carotene, (**B**) pAKm-Zea for zeaxanthin, (**C**) pAKm-Cantha for canthaxanthin, (**D**) pAKm-Asta for astaxanthin. (**E**) The absorbance spectrum and chemical structure of the compound of each major peak. Panels (**F-I**) display mass spectra (top) or product ion spectra (bottom) for the corresponding retinoids: (**F**) retinal, (**G**) 3-hydroxy-retinal (zea-retinal), (**H**) 4-oxo-retinal (cantha-retinal), and (**I**) 3-hydroxy-4-oxo-retinal (asta-retinal). The chemical formula and the calculated exact mass are shown below each compound structure.

The retinoid production distribution showed varied results among the substrates. Zea-retinal (**7**) was produced with lower efficiency compared to cantha-retinal (**8**) and asta-retinal (**9**) (**Fig. 3B-D**). For canthaxanthin and astaxanthin, this study provides the first demonstration of their cleavage by Blh. For zeaxanthin, interestingly, while previous *in vitro* studies^23^ reported no cleavage by Blh, our *in bacterio* results indicate that cleavage does occur, albeit at lower efficiency. This apparent discrepancy may be attributed to differences in reaction conditions between *in vitro* and *in bacterio* (see Discussion).

Another notable observation was the substantial production of retinal and retinol—β-carotene cleavage products. The same carotenoid plasmids, when introduced individually into *E. coli*, result in a single carotenoid (**Fig. 3A-D**, left chromatogram), i.e. not accumulating the intermediate β-carotene. This pattern suggests that Blh more efficiently sequesters β-carotene from the carotenoid pathway compared to the competing enzymes CrtW and CrtZ (see **Fig. 1C**) in *E. coli*. The important consideration here is that the expression level for Blh is considerably higher than that of CrtW/CrtZ: while the *blh* gene was expressed from a high-copy pUC vector (100-300 copies/cell) under a strong IPTG-inducible P_tac_ promoter control, the carotenoid biosynthetic genes were expressed from a medium-copy p15A vector (approx. 20 copies/cell) under a weak promoter (P_J23115_). To modulate the cleavage product distribution, subsequent experiments focused on reducing Blh expression levels and implementing stringent promoter control to optimize expression timing in *E. coli*.

### 4. Tuning Blh expression to modulate retinoid proportions in the engineered pathway

To test whether Blh expression level controls cleavage product distribution in astaxanthin-producing *E. coli*, we constructed four Blh expression variants (**Fig. 4A** and **4B**), varying promoter (P_tac_ vs P_tet_), origin, and RBS strength (weak RBS [Rw]: TIR 135, medium RBS [Rm]: TIR 1878, strong RBS [Rs]: TIR 15549). To assess both expression-level effects and the tightness of induction control, each construct was tested with and without inducer. First, the P_tac_ promoters were tested (**Fig. 4A**). Under full induction, pUC-P_tac_-R_m_-Blh produced predominantly retinal/retinol (∼64%, **Fig. 4A**), and no induction resulted in a similar profile, confirming the well-known leakiness of Ptac. In contrast, the attenuated construct pSC-P_tac_-R_w_-Blh (low-copy origin, weak RBS) under full IPTG induction yielded a more even distribution (∼33% each retinal/retinol, cantha-retinal, and asta-retinal). Without IPTG, this construct showed a further shift toward cantha-retinal (∼48%) and asta-retinal (∼35%), with retinal/retinol reduced to ∼17%. Together, these Ptac data indicate that retinoid product distribution is sensitive to functional Blh activity, but tight expression control could not be achieved with the leaky P_tac_ promoter.

**Figure 4.**
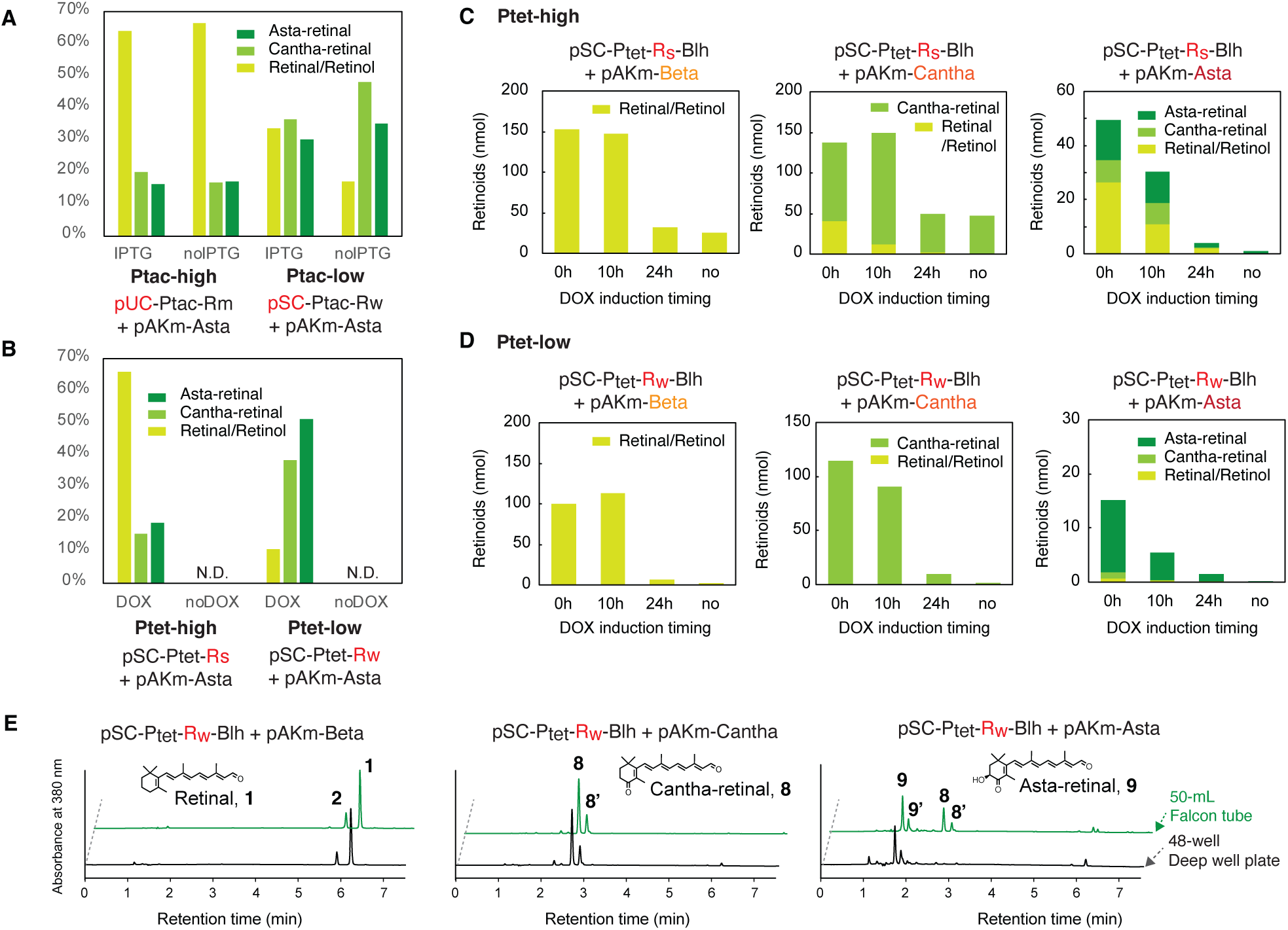
Tuning Blh expression for efficient retinoid production in engineered *E. coli*. (**A, B**) Effect of Blh expression on retinoid product distribution in astaxanthin-producing *E. coli* (harboring pAKm-Asta), comparing P_tac_-driven (**A**) and P_tet_-driven (**B**) Blh constructs. Bars show the molar fraction of each retinoid product; retinal/retinol, cantha-retinal (4-oxo-retinal), and asta-retinal (3-hydroxy-4-oxo-retinal). Calculated from LC-MS peak areas. ’High’ and ’low’ indicate relative functional Blh activity within each promoter family, achieved through different combinations of copy number and RBS strength (see **Methods** and **Supplementary Table 1** for full construct details). Cultures were induced with 1 mM IPTG (**A**) or 100 ng/mL DOX (**B**) at inoculation in 50-mL conical tubes and sampled at 48 h post-induction. (**C, D**) Comparison of induction timing for Blh expression using pSC-P_tet_-R_s_-Blh (strong RBS, **C**) or pSC-P_tet_-R_w_-Blh (weak RBS, **D**), each co-expressed with one of three carotenoid biosynthesis plasmids (pAKm-Beta, pAKm-Cantha, pAKm-Asta). Cultures in 48-well deep well plates were induced with 100 ng/mL DOX at 0, 10 or 24 h post-inoculation, or left uninduced (“no”); retinoids were quantified at 48 h. Stacked bars indicate the molar amount (nmol) of each retinoid product. (**E**) Chromatogram chart of the retinoid production using pSC-P_tet_-R_w_-Blh construct. From left to right, pAKm-Beta + pSC-P_tet_-R_w_-Blh, pAKm-Cantha + pSC-P_tet_-R_w_-Blh, pAKm-Asta + pSC-P_tet_-R_w_-Blh. Top row: 50-mL tube culture. Bottom row: 48-well deep well plate.

Next, to enable higher control, we used a stringent P_tet_ promoter. The constructs were placed on a low copy pSC101 ori, and used either strong RBS (Rs) or weak RBS (Rw) (**Fig. 4B**). Without DOX induction, both P_tet_ constructs produced essentially no retinoids (Fig. 4B, no DOX conditions) confirming stringent regulation. Under DOX induction, pSC-P_tet_-R_s_-Blh (strong RBS) gave a retinal/retinol-dominant profile (∼66%) similar to the high-expression Ptac construct, indicating that strong translational initiation drives premature cleavage of β-carotene intermediates before completion of the astaxanthin pathway. In contrast, pSC-P_tet_-R_w_-Blh (weak RBS) shifted the distribution dramatically: ∼50% asta-retinal and ∼40% cantha-retinal, with only ∼10% retinal/retinol. Together, these results establish that combining the stringent Ptet promoter with weak RBS provides tight control over Blh expression and shifts the cleavage product distribution toward the deeper-derivatized retinoids, with combined cantha-retinal and asta-retinal accounting for ∼90% of total retinoids.

Given the favorable performance of the pSC-P_tet_-R_w_-Blh construct, we next asked two questions: whether this expression-driven control of product distribution extends across different carotenoid substrates, and whether delaying induction (to allow upstream carotenoid biosynthesis to progress further before cleavage) could reduce the premature β-carotene cleavage observed with stronger Blh expression (**Fig. 4C** and **4D**). To address these, *E. coli* harboring either pAKm-Beta, pAKm-Cantha, or pAKm-Asta with either pSC-P_tet_-R_s_-Blh (strong RBS) (**Fig. 4C**) or pSC-P_tet_-R_w_-Blh (weak RBS) (**Fig. 4D**) were induced with 100 ng/mL DOX at 0 h, 10 h and 24 h post-inoculation in 48-well deep well plate cultures, with product analysis performed at 48 h. As a result, two trends emerged. First, the weak-RBS construct (pSC-P_tet_-R_w_-Blh; **Fig. 4D**) at t=0 h induction outperformed all other conditions in producing the downstream cleavage products in pAKm-Cantha and pAKm-Asta. Second, although delayed induction at 10 h with the strong-RBS construct (pSC-P_tet_-R_s_-Blh; **Fig. 4C**) did modestly reduce the retinal/retinol fraction compared to 0 h induction, consistent with our hypothesis that delayed induction allows upstream carotenoid biosynthesis to progress further, the effect was small and substantial retinal/retinol production persisted regardless of timing. By contrast, switching to the weak-RBS construct shifted the distribution dramatically even at 0 h induction. Furthermore, induction at 24 h yielded minimal total product across all conditions, likely reflecting reduced Blh expression in stationary-phase cultures. Together, these results indicate that the relative expression level of Blh, not induction timing, primarily determines product distribution: lowering Blh activity through weak RBS is a more effective lever than delaying its expression. Induction timing, however, significantly affects total retinoid yield, with 0–10 h post-inoculation being optimal.

Having established construct-level control over product distribution through expression tuning, we noticed that the asta-retinal fraction was consistently higher in 48-well plates (**Fig. 4D**) than in 50-mL tube cultures (**Fig. 4B**) using the same construct. To further confirm, we directly compared the product chromatograms across the two culture formats for all three carotenoid substrates (**Fig. 4E**). Across the two culture formats, the chromatograms revealed a substrate-specific pattern. For pAKm-Beta and pAKm-Cantha, the product chromatograms were essentially indistinguishable between 50-mL tube and 48-well plate cultures, indicating that the upstream carotenoid pathways for these substrates are robust to scale-dependent variation. In contrast, for pAKm-Asta, the 50-mL tube culture resulted in a mixture of cantha-retinal and asta-retinal, while the 48-well plate culture showed a cleaner profile dominated by asta-retinal. This difference suggested that the upstream carotenoid pathway reached completion more readily in 48-well plates than in 50-mL tubes. This interpretation is supported by our previous finding that *E. coli* harboring pACm-Asta (differing from pAKm-Asta only in the antibiotic resistance marker) produced almost exclusively astaxanthin at both 24 h and 48 h in 48-well plate cultures, whereas 50-mL tube cultures, despite yielding predominantly astaxanthin by 48 h, accumulated pathway intermediates such as adonixanthin and canthaxanthin at 24 h.^38^

Thus, the substrate-specific scale dependence observed for pAKm-Asta most likely reflects scale/aeration-dependent differences in the longer astaxanthin biosynthetic pathway, which involves multiple ketolation and hydroxylation steps and likely generates different intermediate pool compositions under different culture conditions. This finding has practical implications for scale-up: process re-optimization at each scale will be required to maintain a defined product profile, particularly for substrates with longer or more aerobically demanding biosynthetic pathways. Nonetheless, the expression-level strategy established here provides a robust and transferable framework for controlling retinoid product distribution across different carotenoid substrates.

### 5. Engineering the predicted substrate cavity of Blh for improved zeaxanthin cleavage

Because zeaxanthin cleavage by Blh was inefficient, we examined the enzyme’s predicted structure to investigate possible structural constraints. No crystal structure of Blh or close homologs has been reported. An earlier homology-modeling study, which relied on distant (non-homologous) templates, proposed His-21, His-78, His-188 and His-192 as the four iron-coordinating residues^23^. To re-examine the active site with greater confidence, we analyzed the AlphaFold^39,40^-predicted structure of Blh (UniProt accession Q4PNI0; Blh from Uncultured marine bacterium 66A03), which has an average pLDDT score of 93.68, indicating high confidence in the predicted backbone (**Fig. 5A**). The structure revealed nine transmembrane alpha helices surrounding a distinct internal cavity (detected by pyKVFinder^41^). Within this cavity, three of the four previously identified histidine residues (His-21, 78, 188) lie in a single plane together with His-256, a residue that has not previously been identified as a candidate iron-coordinating residue but is conserved across all 500 Blh/Brp homologs in our alignment (**Fig. 5B; see below**). His-192, by contrast, is positioned outside this plane, above the putative iron-binding site (**Fig. 5B**); this geometry suggests His-192 may not be directly coordinating Fe^2+^, but it may contribute to a second-shell or other auxiliary role. A definitive assignment of the residues will require experimental validation and is beyond the scope of this work; for the analyses that follow, we treat the His-21/78/188/256 plane as the catalytic iron site.

**Figure 5.**
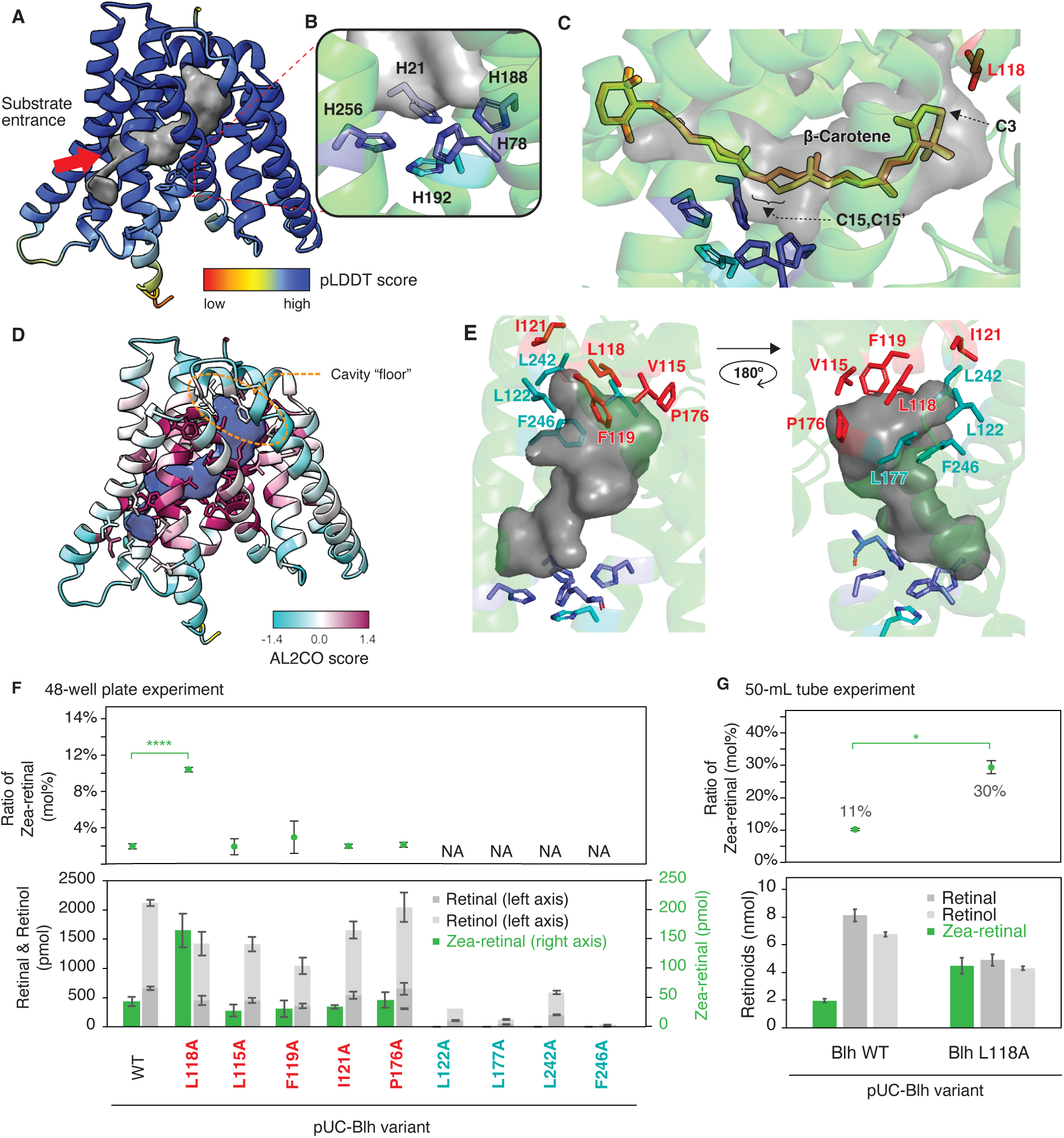
**Predicted structure of Blh, substrate docking and mutagenesis**. (**A**) AlphaFold2 prediction of Blh structure. Retrieved from the AlphaFold Protein Structure Database (UniProt accession: Q4PNI0, AFDB entry: AF-Q4PNI0-F1-v4). The structure is colored by pLDDT score. The substrate cavity was detected by pyKVfinder and shown in grey. (**B**) Close-up of the putative catalytic site showing three previously reported histidine residues (His-21, His-78, His-188) and a newly identified His-256, positioned in the same coordination plane. Previously predicted His-192 (see main text) are indicated as well. (**C**) Docking of β-carotene into the predicted Blh structure using AutoDock Vina.^42,43^ The top two energetically favorable docking poses are shown (indicated in yellow and orange). The 15,15′-cleavage site positioned adjacent to the catalytic site is shown. The C3 position of β-carotene (hydroxy group in zeaxanthin) is indicated. (**D**) Conservation mapping based on an alignment of 140 non-redundant Blh/Brp homologs (reduced from 500 PSI-BLAST hits by CD-HIT at 70% sequence identity), calculated using the entropy-based measure (AL2CO^53^) implemented in ChimeraX; the most highly conserved positions are shown in maroon and the least conserved in cyan. The substrate cavity is shown in blue. (**E**) Close-up view of the cavity floor. Residues targeted for alanine substitution are mapped onto the structure. The 180° -flipped view is indicated on the left. (**F**) Retinoid product analysis screening by Blh alanine-substituted variants (on pUC vector) expressed in *E. coli* harboring pAKm-Zea. Molecular % of zea-retinal (top) and bar graph of retinoid production (bottom). (**G**) Retinoid production in zeaxanthin-producing *E coli* by Blh and Blh-L118A. Molecular % of zea-retinal (top) and bar graph of retinoid production (bottom). For **F** and **G**, bar represents average of 3 replicates and error bars indicate standard deviation. ND, not detected. The structures are shown using ChimeraX or PyMOL.

We performed molecular docking of the natural substrate, β-carotene, into the predicted Blh structure (**Fig. 5C**), using AutoDock Vina^42,43^ (see Methods). The top two energetically favorable docking poses revealed that β-carotene is consistently accommodated within this cavity, with the 15,15’-cleavage site positioned in proximity to the predicted iron site (His-21/78/188/256 plane), consistent with the known cleavage chemistry of Blh.^23^ In all top-scoring poses, C3 (the position bearing a hydroxyl group in zeaxanthin) lies ∼3.6 Å from the Cγ of L118, too close to accommodate a hydroxyl substituent (**Fig. 5C**). In contrast, the cavity provided ample space around C4, the position bearing a keto group in canthaxanthin. This spatial difference may explain the efficient cleavage of canthaxanthin but not zeaxanthin by Blh.

To assess whether this cavity is structurally constrained, we mapped sequence conservation from 140 non-redundant Brp/Blh homologs (reduced from 500 PSI-BLAST hits by CD-HIT at 70% sequence identity) onto the predicted structure (**Methods** and **Fig. 5D**). While overall homology was 10-30%, key residues were highly conserved. Especially, the residues lining the cavity side walls are highly conserved, whereas those forming the cavity floor shows greater variability. This pattern suggests that the cavity floor may be more tolerant to substitutions, making it a plausible target for cavity expansion.

To test whether expanding the cavity floor would enable more efficient zeaxanthin cleavage, we systematically introduced alanine substitutions at residues forming the cavity floor (V115, L118, F119, I121, L122, P176, L177, L242, F246; **Fig. 5E**). Each Blh variant was co-expressed with pAKm-Zea in *E. coli*, and activity was screened in 48-well plate cultures (**Fig. 5F**). Mutations L122A, L177A, L242A, and F246A (indicated in teal in **Fig. 5E**) abolished activity, whereas the remaining five (V115A, L118A, F119A, I121A, P176A; red in **Fig. 5E**) retained it. Among the five active variants, L118A not only retained activity but markedly increased zea-retinal production: while wild-type Blh yielded only ∼2% zea-retinal of total retinoids, L118A increased this to ∼10% (**Fig. 5F**).

To verify this result at a larger scale, we cultured *E. coli* expressing wildtype Blh or L118A together with the zeaxanthin-producing pAKm-Zea construct in 50-mL tube cultures, which use different conditions than the 48-well cultures (see Section 4 and **Fig. 4E**). Consistent with the 48-well screening result, the proportion of zea-retinal increased, from 11% in wild-type to 30% in L118A (**Fig. 5G**). Together with the docking analysis, this result indicates that the L118A substitution alleviates the steric clash with the C3 hydroxyl group, thereby enabling more efficient cleavage of hydroxylated carotenoids such as zeaxanthin.

Cavity-floor substitutions have proven a productive strategy for altering substrate scope in isoprenoid enzymes, applied previously to chain-length extension in farnesyl diphosphate synthase^44^ and phytoene synthase.^45,46^ The L118A substitution identified here demonstrates that cavity-floor engineering is also effective for Blh, and this strategy may enable efficient cleavage of other hydroxylated or bulky carotenoid substrates.

## Discussion

Carotenoid cleavage enzymes play crucial roles in the production of various apocarotenoids, including retinal, the chromophore of microbial rhodopsins.^1,2^ Among them, Brp/Blh-family enzymes have been primarily discussed in the context of bacterial rhodopsin gene clusters, notably in marine bacteria.^26,30^ While many studies have examined the evolutionary/ecological aspects of microbial rhodopsins or Blh^28–33^, or used the Brp/Blh enzyme to provide retinals for heterologous rhodopsin reconstructions,^27,47^ its use in microbial production has been limited to β-carotene-derived retinal/retinol; notably, carotenoids commonly found in marine bacteria, such as canthaxanthin and astaxanthin, have never been tested as substrates for retinoid production. In this study, we systematically expressed Blh in an *E. coli* co-expression system and discovered unexpected promiscuous activity toward various carotenoids, and leveraged this promiscuity to produce diverse retinal analogues. Furthermore, we demonstrated that the distribution of retinoid products could be modulated through careful regulation of Blh expression and rational enzyme engineering.

Our findings both corroborate and extend previous knowledge of Blh’s substrate specificity. Kim et al.^23^ conducted the first comprehensive *in vitro* analysis of Blh activity using a soluble fraction of the purified enzyme, despite its predicted membrane protein nature. Their study reported activity toward β-carotene, β-cryptoxanthin, α-carotene, γ-carotene, and β-apo-4’-carotenal, while zeaxanthin, lutein, lycopene, and shorter apocarotenoids remained un-cleaved. Our *in bacterio* results confirmed the efficient cleavage of β-carotene and γ-carotene, yielding their respective retinals and alcohol derivatives. Interestingly, we observed trace cleavage of zeaxanthin to zea-retinal, which was not detected in previous studies. While this could potentially result from the cleavage of β-cryptoxanthin transiently accumulated during zeaxanthin biosynthesis, our engineered *E. coli* expression system consistently produced small amounts of zea-retinal, providing a platform for further investigation. It is possible that our intracellular expression system allowed us to assess Blh’s ability to cleave a range of carotenoids within a living cellular environment, thereby bypassing the challenges of *in vitro* purification and potentially overcoming limitations observed in previous studies.

A significant finding of our studies was the identification of novel Blh substrates. The enzyme exhibited activity toward δ-carotene (ε-ring, ψ-end group) and ε-carotene (ε,ε-rings), generating α-retinal (ε-ring). While the previous study^23^ reported α-carotene (ε,β-rings) cleavage, the precise ring specificity remained unclear. Our results demonstrate that Blh can cleave at the ε-ring, albeit with reduced efficiency compared to β-ring substrates. This promiscuous activity is particularly intriguing as ε-ring carotenoids are predominantly found in plants, algae and eukaryotes, with bacterial occurrence limited to few genera of cyanobacteria.^48^ The observed activity might represent inherent promiscuity toward structures rarely encountered in Blh’s natural environment. Most notably, we discovered that wild-type Blh efficiently cleaves canthaxanthin and astaxanthin, generating cantha-retinal and asta-retinal, respectively. Especially, the amount of cantha-retinal produced was comparable to that of retinal (**Fig. 3C**).

The range of carotenoids accepted by Blh raises intriguing questions about its biological roles. Canthaxanthin and astaxanthin are frequently found in marine bacteria, suggesting potential co-occurrence with bacterial rhodopsin systems. Here, we performed a homology search using phmmer^49^ against Ensembl bacteria database (via the web server https://bacteria.ensembl.org/hmmer/index.html) using *blh* used in this study and *crtW* (β-carotene ketolase gene) from *Brevundimonas* sp. SD212 as queries. Using an E-value cutoff 1e-5 (a measure of statistical significance), we identified 13 organisms containing both genes. By increasing the E-value cutoff to allow for less stringent matches, the hit expanded to 32 organisms. Given that our analysis used single representative sequences as queries, the actual co-occurrence is likely higher. The conserved residue mapped onto the Blh predicted structure (**Fig. 5D**) also shows that the cavity floor is not highly conserved. Our results, demonstrating that single substitutions in this region led to shifted substrate specificity, suggest that acquiring new functions may be relatively straightforward. Some Blh homologs might have evolved to utilize alternative carotenoids, producing diverse retinoids with distinct biological functions. This hypothesis is particularly compelling given the growing recognition of varied physiological roles for retinoids and apocarotenoids in biological systems, and the recent discovery of Blh homologs in many eukaryotes.^28^ Conversely, if most of Blh homologs exhibit high specificity solely towards β-carotene, this would suggest a strong evolutionary pressure towards that specificity, which also would be an interesting finding.

Our *E. coli* co-expression system provides a robust platform for rapid characterization of these enzymes’ promiscuity, potentially unveiling unexpected diversity in their biological functions. Furthermore, the demonstrated substrate promiscuity of Blh opens possibilities for generating novel retinoid structures. Given the diverse biological activities of retinoids across different organisms, engineered Blh variants might enable the production of new bioactive compounds with potential therapeutic or biotechnological applications. The findings in this study will be particularly useful to biochemists interested in the reaction mechanism of Blh. Further investigation of natural Blh diversity may reveal additional capabilities and applications of these versatile enzymes.

## Materials and methods

### Strains and reagents

*E. coli* strains used in this study were either NEB Turbo (New England BioLabs, Ipswich, MA) or DH5α (SMOBIO Technology, Inc., Taiwan). *E. coli* was cultured in LB-Lennox (Nacalai Tesque, Inc., Kyoto, #20066-24) or Terrific Broth (TB) media (tryptone 12 g, yeast extract 24 g, glycerol 4 mL, KH_2_PO_4_ 2.323 g, K_2_HPO_4_ 12.54 g per 1 L water). Antibiotics used were chloramphenicol (FUJIFILM Wako, Osaka, #036-10571), carbenicillin (Wako #594-02193) or kanamycin (Wako #117-00343) with the final concentration of 30, 50 or 50 µg/mL, respectively. Inducers used were isopropyl β-D-thiogalactopyranoside (IPTG) (Wako #096-05143) and doxycycline (DOX) (Wako #049-31121). As for the authentic standard for LC-PDA-MS, retinyl acetate (Wako #184-02891), all-trans-retinal (Wako #180-03111), and all-trans-retinol (Wako #R1876) were purchased.

### Plasmid construction

Plasmids used in this study are listed in **Supplementary Table 1**. The carotenoid plasmids except for pAKm-Gamma were derived from the previous study.^38^ pAKm-Gamma was constructed as follows: pUClac-crtYcYd was constructed by amplifying the *crtYc-crtYd* genes from *Myxococcus xanthus*^50^ using crtYcYd-F/R primers (**Supplementary Table 2**) and inserting into the pUClac vector^51^ by homology cloning. The *crtYcYd* fragment was amplified using oMF361/362 primers and inserted into the XhoI site of pACm-Lyco^38^ by homology assembly.

Plasmids for Blh expression were constructed as follows. The protein sequence of Blh from Uncultured marine bacterium 66A03^23^ was obtained from Uniprot database (accession no. Q4PNI0). The *E. coli* codon-optimized sequence (**Supplementary Table 2**) was designed and a strong ribosome binding site (RBS) sequence (5’-AGGAGGATTACAA-3’) was added to the upstream. The ordered DNA fragment (gBlock gene fragments, IDT DNA, IA, USA) was inserted in XhoI/HindIII site of pUClac vector^51^ using homology cloning and named this plasmid as pUC-P_tac_-R_s_-Blh (R_s_ as strong RBS). A weaker RBS with translation initiation rate (TIR) of 100 or 2000 was designed by RBS calculator^37,52^ from De Novo DNA (www.denovodna.com). The calculated sequence, weak RBS sequence (5’-AGATATAATCGAACCAT −3’) with TIR 135, and medium RBS sequence (5’-AGCGAGGGGGTTTCCAT −3’) with TIR 1878, was replaced with the original strong RBS sequence, and the construct was named pUC-P_tac_-R_w_-Blh and pUC-P_tac_-R_m_-Blh.

pSC-P_tac_-R_w_-Blh and pSC-P_tac_-R_s_-Blh were constructed by amplifying the Blh fragment by oMFseq45/oMFseq47 and digesting with XhoI/HindIII, and ligating into XhoI/HindIII-cleaved pSCAp1.1e vector (pMF821): this vector was constructed by combining the fragments of pSC101 ori, P_tac_/*lacI*-*lacZ* and Amp^R^ marker (sequence provided in **Supplementary Table 2**). pSC-P_tet_-R_w_-Blh and pSC-P_tet_-R_s_-Blh were constructed by cleaving out the P_tac_/lacI fragment by PvuI/XhoI sites and inserting a gene fragment of P_LtetO-1_/tetR fragment by homology cloning.

### Bacterial culture

The *E. coli* culture was performed at two scales: in 50-mL falcon tube (**Fig. 2, 3, 4A, 4B, 4E, 5G**) or in 48-well deep well plate (**Fig. 4C-E** and **5F**). For the 50-mL falcon tube culture, the transformant *E. coli* colonies carrying the plasmids were picked and inoculated into 2 mL LB-media supplemented with antibiotics in 2-position round bottom tube and cultured at 37°C, 200 rpm overnight. A portion (50 µL) of culture was transferred to 10-mL TB-media supplemented with antibiotics/inducers in a 50-mL falcon tube. Dodecane (1 mL) was added on top of the inoculated media. The cells were cultured at 30°C, 200 rpm for 48 hours.

For the 48-well plate culture, the transformant *E. coli* colonies carrying the plasmids were picked and inoculated into 1 mL LB-media supplemented with antibiotics in 48-well deep well plate covered with porous film seal (AeraSeal, Excel Scientific, Inc.) and cultured at 37°C, 1000 rpm overnight. A portion (20 µL) of culture was transferred to 1-mL TB-media supplemented with antibiotics/inducers in a 48-well deep well plate. Dodecane (200 µL or 500 µL) was added on top of the inoculated media. The plate was covered with AeraSeal and the cells were cultured at 30°C, 1000 rpm for 48 hours. Inducer (DOX) was added with a final concentration of 100 ng/mL.

### Retinoid and carotenoid extraction

For the 50-mL culture, the culture tubes were centrifuged at 8000*g* for 3 min, and the dodecane overlays (approx. 750 µL) was collected using pipette to a glass vial. The cells were re-harvested by centrifuged at 8000*g* for 3 min. The supernatant was discarded, and the pellet was washed with 5 mL 1% NaCl solution. After discarding the supernatant, the pellets were vortexed for approximately 5 min until they become soft. Five mL of acetone was added to the cell pellet and vortexed immediately for 5 min. The tubes were centrifuged at max speed for 5 min and the acetone extracts were collected.

For the 48-well deep well plate culture, the cell cultures (together with dodecane layer) were collected with pipette to a new 2-mL collection tube. The tubes were centrifuged at 8000g for 3 min, and the top dodecane overlays were collected to new collection tubes by pipette (approx. 200 µL). The tubes were centrifuged again, and the supernatant was discarded, and the 1-mL 1% NaCl solution was added followed by vortex. After centrifugation, the supernatant was discarded, and the pellets were vortexed for approximately 5 min until they become soft. Acetone (500 µL) was added to the pellet and vortexed for 5 min. The tubes were centrifuged at max speed for 2 min and the acetone extracts were collected.

### Retinoid and carotenoid analysis

The dodecane phases or acetone extracts were analyzed with liquid chromatography equipped with photodiode array and mass spectrometry (LC-PDA-MS). The equipment used were Waters UPLC H-Class PLUS consisting of a quaternary pump with degasser, autosampler and photodiode array (PDA) in combination with a Xevo TQ-S micro triple quadrupole mass spectrometer. The column used was Waters ACQUITY UPLC BEH C18 1.7 µm column (2.1 mm x 100 mm) equipped with a C18 VanGuard Pre-column. Injection volumes of the samples were 5 µL for dodecane phases and 2 or 2.5 µL for acetone extracts. The mobile phase used were A: water/acetonitrile = 30/70, B: acetonitrile, C: methanol, D: isopropanol with an elution flux of 0.2 ml/min with the following gradient program (total 15 min). Starting from [100% A], 0-2 min gradient to [70% A, 30% B]; 2-4 min gradient to [35% B, 15% C, 50% D] and hold for 8 min until 12 min; switch to [100% A] and hold for 3 min. The elution was monitored with absorbance at 450 nm (for carotenoid acetone extracts) or the max plot of 300-400 nm (for dodecane phases). The data was analyzed using Waters MassLynx software.

### Sequence Alignment of Blh homologs and structural mapping

Using Blh from Uncultured marine bacterium 66A03^23^ (UniProt accession no. Q4PNI0) as a query, PSI-BLAST was performed against the NCBI nr database (accessed 2025/4/26), identifying 500 Brp/Blh homologs (a threshold E-value of 3e-42). Redundant sequences were removed with CD-HIT (v.4.8.1) using a sequence identity threshold of 70%, yielding 140 non-redundant Brp/Blh homologs. The sequences were aligned with MAFFT (v7.490) using Geneious (Algorithm: Auto, Scoring matrix: BLOSUM62, Gap open penalty: 1.53, Offset value: 0.123). The resulting alignment was mapped onto the AlphaFold2 model of Blh in ChimeraX (v.1.9), which assigns per-residue conservation values (the “seq_conservation” attribute) calculated using the entropy-based measure from AL2CO.^53^

### Substrate docking of Blh and β-carotene

The docking of AlphaFold2 structure of Blh and β-carotene substrate was performed using AutoDock Vina.^42,43^ The coordinate of the Blh protein was obtained using PyMOL cmd.get_coords command, and the β-carotene substrate was obtained from PubChem (CID 5280489). The ligand and receptor PDBQT files were prepared with Meeko.^54^ Docking was performed with the receptor treated as rigid, using a search box of 30 x 30 x 30 Å centered at (1.99, 0.77, 2.42) and an exhaustiveness of 32. The two most energetically favorable docking poses with affinity of −10.9 kcal/mol were selected, and the structure was visualized in ChimeraX (v1.9) or PyMOL (v2.5.0).

## Data availability

All data generated or analyzed during this study are included in this published article and its supplementary information file.

## Acknowledgements

M.F. thanks M. Nariai, M. Hayakawa and Y. Miyamura for the technical support, and Dr. Yoshiaki Yasutake (AIST) for the discussion on molecular docking. This study is funded by KAKENHI 23H02144 and 23K26837.

## References

(1) Kandori, H. Biophysics of Rhodopsins and Optogenetics. Biophys Rev 2020, 12 (2), 355–361. 10.1007/s12551-020-00645-0.

(2) Rozenberg, A.; Inoue, K.; Kandori, H.; Béjà, O. Microbial Rhodopsins: The Last Two Decades. Annu Rev Microbiol 2021, 75, 427–447. 10.1146/annurev-micro-031721-020452.

(3) Das, B. C.; Thapa, P.; Karki, R.; Das, S.; Mahapatra, S.; Liu, T.-C.; Torregroza, I.; Wallace, D. P.; Kambhampati, S.; Van Veldhuizen, P.; Verma, A.; Ray, S. K.; Evans, T. Retinoic Acid Signaling Pathways in Development and Diseases. Bioorganic & Medicinal Chemistry 2014, 22 (2), 673–683. 10.1016/j.bmc.2013.11.025.

(4) Álvarez, R.; Vaz, B.; Gronemeyer, H.; de Lera, Á. R. Functions, Therapeutic Applications, and Synthesis of Retinoids and Carotenoids. Chem. Rev. 2014, 114 (1), 1–125. 10.1021/cr400126u.

(5) Liang, C.; Qiao, G.; Liu, Y.; Tian, L.; Hui, N.; Li, J.; Ma, Y.; Li, H.; Zhao, Q.; Cao, W.; Liu, H.; Ren, X. Overview of All-Trans-Retinoic Acid (ATRA) and Its Analogues: Structures, Activities, and Mechanisms in Acute Promyelocytic Leukaemia. European Journal of Medicinal Chemistry 2021, 220, 113451. 10.1016/j.ejmech.2021.113451.

(6) Jang, H.-J.; Yoon, S.-H.; Ryu, H.-K.; Kim, J.-H.; Wang, C.-L.; Kim, J.-Y.; Oh, D.-K.; Kim, S.-W. Retinoid Production Using Metabolically Engineered Escherichia Coli with a Two-Phase Culture System. Microbial Cell Factories 2011, 10 (1), 59. 10.1186/1475-2859-10-59.

(7) Jang, H.-J.; Ha, B.-K.; Zhou, J.; Ahn, J.; Yoon, S.-H.; Kim, S.-W. Selective Retinol Production by Modulating the Composition of Retinoids from Metabolically Engineered E. Coli. Biotechnology and Bioengineering 2015, 112 (8), 1604–1612. 10.1002/bit.25577.

(8) Han, M.; Lee, P. C. Microbial Production of Bioactive Retinoic Acid Using Metabolically Engineered Escherichia Coli. Microorganisms 2021, 9 (7), 1520. 10.3390/microorganisms9071520.

(9) Hu, Q.; Yu, H.; Ye, L. Production of Retinoic Acid by Engineered Saccharomyces Cerevisiae Using an Endogenous Aldehyde Dehydrogenase. Biotechnology and Bioengineering 2022, 119 (11), 3241–3251. 10.1002/bit.28192.

(10) Wang, X.; Xu, X.; Liu, J.; Liu, Y.; Li, J.; Du, G.; Lv, X.; Liu, L. Metabolic Engineering of Saccharomyces Cerevisiae for Efficient Retinol Synthesis. Journal of Fungi 2023, 9 (5), 512. 10.3390/jof9050512.

(11) Shi, Y.; Lu, S.; Zhou, X.; Wang, X.; Zhang, C.; Wu, N.; Dong, T.; Xing, S.; Wang, Y.; Xiao, W.; Yao, M. Systematic Metabolic Engineering Enables Highly Efficient Production of Vitamin A in *Saccharomyces Cerevisiae*. Synthetic and Systems Biotechnology 2025, 10 (1), 58–67. 10.1016/j.synbio.2024.08.004.

(12) Shin, K.-C.; Seo, M.-J.; Kim, Y.-S.; Yeom, S.-J. Molecular Properties of β-Carotene Oxygenases and Their Potential in Industrial Production of Vitamin A and Its Derivatives. Antioxidants (Basel) 2022, 11 (6), 1180. 10.3390/antiox11061180.

(13) Beltran, J. C. M.; Stange, C. Apocarotenoids: A New Carotenoid-Derived Pathway. In Carotenoids in Nature: Biosynthesis, Regulation and Function; Stange, C., Ed.; Subcellular Biochemistry; Springer International Publishing: Cham, 2016; pp 239–272. 10.1007/978-3-319-39126-7_9.

(14) Harrison, E. H.; Quadro, L. Apocarotenoids: Emerging Roles in Mammals. Annu. Rev. Nutr. 2018, 38 (1), 153–172. 10.1146/annurev-nutr-082117-051841.

(15) Moreno, J. C.; Mi, J.; Alagoz, Y.; Al-Babili, S. Plant Apocarotenoids: From Retrograde Signaling to Interspecific Communication. The Plant Journal 2021, 105 (2), 351–375. 10.1111/tpj.15102.

(16) Daruwalla, A.; Kiser, P. D. Structural and Mechanistic Aspects of Carotenoid Cleavage Dioxygenases (CCDs). Biochimica et Biophysica Acta (BBA) - Molecular and Cell Biology of Lipids 2020, 1865 (11), 158590. 10.1016/j.bbalip.2019.158590.

(17) Seña, C. dela; Riedl, K. M.; Narayanasamy, S.; Curley, R. W.; Schwartz, S. J.; Harrison, E. H. The Human Enzyme That Converts Dietary Provitamin A Carotenoids to Vitamin A Is a Dioxygenase *. Journal of Biological Chemistry 2014, 289 (19), 13661–13666. 10.1074/jbc.M114.557710.

(18) von Lintig, J.; Vogt, K. Vitamin A Formation in Animals: Molecular Identification and Functional Characterization of Carotene Cleaving Enzymes. J Nutr 2004, 134 (1), 251S–256S. 10.1093/jn/134.1.251S.

(19) Poliakov, E.; Uppal, S.; Rogozin, I. B.; Gentleman, S.; Redmond, T. M. Evolutionary Aspects and Enzymology of Metazoan Carotenoid Cleavage Oxygenases. Biochimica et Biophysica Acta (BBA) - Molecular and Cell Biology of Lipids 2020, 1865 (11), 158665. 10.1016/j.bbalip.2020.158665.

(20) Ahrazem, O.; Gómez-Gómez, L.; Rodrigo, M. J.; Avalos, J.; Limón, M. C. Carotenoid Cleavage Oxygenases from Microbes and Photosynthetic Organisms: Features and Functions. International Journal of Molecular Sciences 2016, 17 (11), 1781. 10.3390/ijms17111781.

(21) Daruwalla, A.; Zhang, J.; Lee, H. J.; Khadka, N.; Farquhar, E. R.; Shi, W.; Lintig, J. von; Kiser, P. D. Structural Basis for Carotenoid Cleavage by an Archaeal Carotenoid Dioxygenase. PNAS 2020, 117 (33), 19914–19925. 10.1073/pnas.2004116117.

(22) Seña, C. dela; Sun, J.; Narayanasamy, S.; Riedl, K. M.; Yuan, Y.; Curley, R. W.; Schwartz, S. J.; Harrison, E. H. Substrate Specificity of Purified Recombinant Chicken β-Carotene 9′,10′-Oxygenase (BCO2). J. Biol. Chem. 2016, 291 (28), 14609–14619. 10.1074/jbc.M116.723684.

(23) Kim, Y.-S.; Kim, N.-H.; Yeom, S.-J.; Kim, S.-W.; Oh, D.-K. In Vitro Characterization of a Recombinant Blh Protein from an Uncultured Marine Bacterium as a β-Carotene 15,15′-Dioxygenase. J. Biol. Chem. 2009, 284 (23), 15781–15793. 10.1074/jbc.M109.002618.

(24) Betlach, M.; Friedman, J.; Boyer, H. W.; Pfeifer, F. Characterization of a Halobacterial Gene Affecting Bacterio-Opsin Gene Expression. Nucleic Acids Res 1984, 12 (20), 7949–7959. 10.1093/nar/12.20.7949.

(25) Peck, R. F.; Echavarri-Erasun, C.; Johnson, E. A.; Ng, W. V.; Kennedy, S. P.; Hood, L.; DasSarma, S.; Krebs, M. P. Brp and Blh Are Required for Synthesis of the Retinal Cofactor of Bacteriorhodopsin in Halobacterium Salinarum. J. Biol. Chem. 2001, 276 (8), 5739–5744. 10.1074/jbc.M009492200.

(26) Sabehi, G.; Loy, A.; Jung, K.-H.; Partha, R.; Spudich, J. L.; Isaacson, T.; Hirschberg, J.; Wagner, M.; Béjà, O. New Insights into Metabolic Properties of Marine Bacteria Encoding Proteorhodopsins. PLOS Biology 2005, 3 (8), e273. 10.1371/journal.pbio.0030273.

(27) Martinez, A.; Bradley, A. S.; Waldbauer, J. R.; Summons, R. E.; DeLong, E. F. Proteorhodopsin Photosystem Gene Expression Enables Photophosphorylation in a Heterologous Host. Proc Natl Acad Sci U S A 2007, 104 (13), 5590–5595. 10.1073/pnas.0611470104.

(28) Rius, M.; Rest, J. S.; Filloramo, G. V.; Novák Vanclová, A. M. G.; Archibald, J. M.; Collier, J. L. Horizontal Gene Transfer and Fusion Spread Carotenogenesis Among Diverse Heterotrophic Protists. Genome Biology and Evolution 2023, 15 (3), evad029. 10.1093/gbe/evad029.

(29) Klassen, J. L. Phylogenetic and Evolutionary Patterns in Microbial Carotenoid Biosynthesis Are Revealed by Comparative Genomics. PLOS ONE 2010, 5 (6), e11257. 10.1371/journal.pone.0011257.

(30) Nakajima, Y.; Tsukamoto, T.; Kumagai, Y.; Ogura, Y.; Hayashi, T.; Song, J.; Kikukawa, T.; Demura, M.; Kogure, K.; Sudo, Y.; Yoshizawa, S. Presence of a Haloarchaeal Halorhodopsin-Like Cl− Pump in Marine Bacteria. Microbes Environ 2018, 33 (1), 89–97. 10.1264/jsme2.ME17197.

(31) Olson, D. K.; Yoshizawa, S.; Boeuf, D.; Iwasaki, W.; DeLong, E. F. Proteorhodopsin Variability and Distribution in the North Pacific Subtropical Gyre. The ISME Journal 2018, 12 (4), 1047–1060. 10.1038/s41396-018-0074-4.

(32) Nakajima, Y.; Kojima, K.; Kashiyama, Y.; Doi, S.; Nakai, R.; Sudo, Y.; Kogure, K.; Yoshizawa, S. Bacterium Lacking a Known Gene for Retinal Biosynthesis Constructs Functional Rhodopsins. Microbes and Environments 2020, 35 (4), ME20085. 10.1264/jsme2.ME20085.

(33) Yoshizawa, S.; Kumagai, Y.; Kim, H.; Ogura, Y.; Hayashi, T.; Iwasaki, W.; DeLong, E. F.; Kogure, K. Functional Characterization of Flavobacteria Rhodopsins Reveals a Unique Class of Light-Driven Chloride Pump in Bacteria. Proceedings of the National Academy of Sciences 2014, 111 (18), 6732–6737. 10.1073/pnas.1403051111.

(34) Kim, Y.-S.; Oh, D.-K. Biotransformation of Carotenoids to Retinal by Carotenoid 15,15′-Oxygenase. Appl Microbiol Biotechnol 2010, 88 (4), 807–816. 10.1007/s00253-010-2823-9.

(35) Zhang, C.; Chen, X.; Lindley, N. D.; Too, H.-P. A “Plug-n-Play” Modular Metabolic System for the Production of Apocarotenoids. Biotechnology and Bioengineering 2018, 115 (1), 174–183. 10.1002/bit.26462.

(36) Dwulit-Smith, J. R.; Hamilton, J. J.; Stevenson, D. M.; He, S.; Oyserman, B. O.; Moya-Flores, F.; Garcia, S. L.; Amador-Noguez, D.; McMahon, K. D.; Forest, K. T. acI Actinobacteria Assemble a Functional Actinorhodopsin with Natively Synthesized Retinal. Appl. Environ. Microbiol. 2018, 84 (24). 10.1128/AEM.01678-18.

(37) Salis, H. M. The Ribosome Binding Site Calculator. Methods Enzymol 2011, 498, 19–42. 10.1016/B978-0-12-385120-8.00002-4.

(38) Furubayashi, M. Systematic Plasmid Engineering for Targeted Carotenoid Synthesis in Bacteria. bioRxiv December 24, 2024, p 2024.12.23.629938. 10.1101/2024.12.23.629938.

(39) Jumper, J.; Evans, R.; Pritzel, A.; Green, T.; Figurnov, M.; Ronneberger, O.; Tunyasuvunakool, K.; Bates, R.; Žídek, A.; Potapenko, A.; Bridgland, A.; Meyer, C.; Kohl, S. A. A.; Ballard, A. J.; Cowie, A.; Romera-Paredes, B.; Nikolov, S.; Jain, R.; Adler, J.; Back, T.; Petersen, S.; Reiman, D.; Clancy, E.; Zielinski, M.; Steinegger, M.; Pacholska, M.; Berghammer, T.; Bodenstein, S.; Silver, D.; Vinyals, O.; Senior, A. W.; Kavukcuoglu, K.; Kohli, P.; Hassabis, D. Highly Accurate Protein Structure Prediction with AlphaFold. Nature 2021, 596 (7873), 583–589. 10.1038/s41586-021-03819-2.

(40) Varadi, M.; Anyango, S.; Deshpande, M.; Nair, S.; Natassia, C.; Yordanova, G.; Yuan, D.; Stroe, O.; Wood, G.; Laydon, A.; Žídek, A.; Green, T.; Tunyasuvunakool, K.; Petersen, S.; Jumper, J.; Clancy, E.; Green, R.; Vora, A.; Lutfi, M.; Figurnov, M.; Cowie, A.; Hobbs, N.; Kohli, P.; Kleywegt, G.; Birney, E.; Hassabis, D.; Velankar, S. AlphaFold Protein Structure Database: Massively Expanding the Structural Coverage of Protein-Sequence Space with High-Accuracy Models. Nucleic Acids Res 2022, 50 (D1), D439–D444. 10.1093/nar/gkab1061.

(41) Guerra, J. V. da S.; Ribeiro-Filho, H. V.; Jara, G. E.; Bortot, L. O.; Pereira, J. G. de C.; Lopes-de-Oliveira, P. S. pyKVFinder: An Efficient and Integrable Python Package for Biomolecular Cavity Detection and Characterization in Data Science. BMC Bioinformatics 2021, 22, 607. 10.1186/s12859-021-04519-4.

(42) Trott, O.; Olson, A. J. AutoDock Vina: Improving the Speed and Accuracy of Docking with a New Scoring Function, Efficient Optimization and Multithreading. J Comput Chem 2010, 31 (2), 455–461. 10.1002/jcc.21334.

(43) Eberhardt, J.; Santos-Martins, D.; Tillack, A. F.; Forli, S. AutoDock Vina 1.2.0: New Docking Methods, Expanded Force Field, and Python Bindings. J Chem Inf Model 2021, 61 (8), 3891–3898. 10.1021/acs.jcim.1c00203.

(44) Ohnuma, S.; Narita, K.; Nakazawa, T.; Ishida, C.; Takeuchi, Y.; Ohto, C.; Nishino, T. A Role of the Amino Acid Residue Located on the Fifth Position before the First Aspartate-Rich Motif of Farnesyl Diphosphate Synthase on Determination of the Final Product*. Journal of Biological Chemistry 1996, 271 (48), 30748–30754. 10.1074/jbc.271.48.30748.

(45) Furubayashi, M.; Ikezumi, M.; Takaichi, S.; Maoka, T.; Hemmi, H.; Ogawa, T.; Saito, K.; Tobias, A. V.; Umeno, D. A Highly Selective Biosynthetic Pathway to Non-Natural C50 Carotenoids Assembled from Moderately Selective Enzymes. Nature Communications 2015, 6, 7534. 10.1038/ncomms8534.

(46) Li, L.; Furubayashi, M.; Hosoi, T.; Seki, T.; Otani, Y.; Kawai-Noma, S.; Saito, K.; Umeno, D. Construction of a Nonnatural C60 Carotenoid Biosynthetic Pathway. ACS synthetic biology 2019, 8 (3), 511–520.

(47) Kim, S. Y.; Waschuk, S. A.; Brown, L. S.; Jung, K.-H. Screening and Characterization of Proteorhodopsin Color-Tuning Mutations in *Escherichia Coli* with Endogenous Retinal Synthesis. Biochimica et Biophysica Acta (BBA) - Bioenergetics 2008, 1777 (6), 504–513. 10.1016/j.bbabio.2008.03.010.

(48) Takaichi, S.; Mochimaru, M.; Uchida, H.; Murakami, A.; Hirose, E.; Maoka, T.; Tsuchiya, T.; Mimuro, M. Opposite Chilarity of α-Carotene in Unusual Cyanobacteria with Unique Chlorophylls, Acaryochloris and Prochlorococcus. Plant and Cell Physiology 2012, 53 (11), 1881–1888. 10.1093/pcp/pcs126.

(49) Potter, S. C.; Luciani, A.; Eddy, S. R.; Park, Y.; Lopez, R.; Finn, R. D. HMMER Web Server: 2018 Update. Nucleic Acids Res 2018, 46 (W1), W200–W204. 10.1093/nar/gky448.

(50) Iniesta, A. A.; Cervantes, M.; Murillo, F. J. Conversion of the Lycopene Monocyclase of Myxococcus Xanthus into a Bicyclase. Appl. Microbiol. Biotechnol. 2008, 79 (5), 793–802. 10.1007/s00253-008-1481-7.

(51) Furubayashi, M.; Kubo, A.; Takemura, M.; Otani, Y.; Maoka, T.; Terada, Y.; Yaoi, K.; Ohdan, K.; Misawa, N.; Mitani, Y. Capsanthin Production in Escherichia Coli by Overexpression of Capsanthin/Capsorubin Synthase from Capsicum Annuum. J. Agric. Food Chem. 2021, 69 (17), 5076–5085. 10.1021/acs.jafc.1c00083.

(52) Roots, C. T.; Lukasiewicz, A.; Barrick, J. E. OSTIR: Open Source Translation Initiation Rate Prediction. Journal of open source software 2021, 6 (64), 3362. 10.21105/joss.03362.

(53) Pei, J.; Grishin, N. V. AL2CO: Calculation of Positional Conservation in a Protein Sequence Alignment. Bioinformatics 2001, 17 (8), 700–712. 10.1093/bioinformatics/17.8.700.

(54) Santos-Martins, D.; He, Y.; Eberhardt, J.; Sharma, P.; Bruciaferri, N.; Holcomb, M.; Llanos, M. A.; Hansel-Harris, A.; Barkdull, A. P.; Tillack, A. F.; Bianco, G.; Paulsen, M.-L.; Mato, J.; Taneja, I.; Forli, S. Meeko: Molecule Parametrization and Software Interoperability for Docking and Beyond. J. Chem. Inf. Model. 2025, 65 (24), 13045–13050. 10.1021/acs.jcim.5c02271.

