## Supplementary Information for "Production of diverse retinal analogues in engineered *Escherichia coli* through promiscuous carotenoid cleavage by Blh"

**Supplementary Table 1.** Plasmid used in this study.

| Plasmid name | Vector Backbone | Contents | Source |
|---|---|---|---|
| pACm-Beta | p15A, Cm | β-Carotene operon | Ref. 38 |
| pAKm-Beta | p15A, Km | β-Carotene operon | Ref. 38 |
| pAKm-Cantha | p15A, Km | Canthaxanthin operon | Ref. 38 |
| pAKm-Zea | p15A, Km | Zeaxanthin operon | Ref. 38 |
| pAKm-Asta | p15A, Km | Astaxanthin operon | Ref. 38 |
| pAKm-Gamma | p15A, Km | γ-Carotene operon | This study |
| pACm-Lyco | p15A, Cm | Lycopene operon | Ref. 38 |
| pACm-Delta | p15A, Cm | δ-Carotene operon | Ref. 38 |
| pACm-Epsilon | p15A, Cm | ε-Carotene operon | Ref. 38 |
| pUC-$P_{tac}$-gfp | pUC, Amp | $P_{tac}$-gfp | Ref. 51 |
| pUC-$P_{tac}$-Rs-Blh | pUC, Amp | $P_{tac}$-$RBS_{15549}$-Blh | This study |
| pUC-$P_{tac}$-Rm-Blh | pUC, Amp | $P_{tac}$-$RBS_{1878}$ – Blh | This study |
| pUC-$P_{tac}$-Rw-Blh | pUC, Amp | $P_{tac}$-$RBS_{100}$ – Blh | This study |
| pSC-$P_{tac}$-Rw-Blh | pSC101, Amp | $P_{tac}$-$RBS_{100}$ – Blh | This study |
| pSC-$P_{tet}$-Rs-Blh | pSC101, Amp | $P_{tet}$-$RBS_{15549}$ – Blh | This study |
| pSC-$P_{tet}$-Rw-Blh | pSC101, Amp | $P_{tet}$-$RBS_{100}$ – Blh | This study |
| pSCAp1.1e | pSC101, Amp | pSC empty vector | This study (Supplementary Table 2) |

**Supplementary Table 2. DNA sequences**

| | Sequence |
|---|---|
| Blh<br><br>RBS (strong) in red. See below for weak/medium RBS.<br>ORF in blue<br>**XhoI/HindIII in bold** | **ctcgag**gaggattacaaatgggattaatgctaatagattggtgtgctttagcgctggtggtgttcatcgg<br>tttgccacacggcgcactggatgcagcgatctcatttagcatgatcagcagtgccaaacgcatcgcccgt<br>ctggccggtatttttgcttatttatctgctattggcgaccgcatttttttctcatctggtatcaactgccgg<br>ctttctcgctgttgatcttcctgttaatctccatcatccatttcggtatggcagacttcaacgcgtcccc<br>gagcaagctgaagtggccgcacattattgcgcatggtggggtggttaccgtttggctgccgctcattcag<br>aaaaacgaagtaacgaaactgttctctattttgactaacggtccgaccccgattctgtgggacatcctgc<br>tgatctttttcctgtgttggagcatcggcgtgtgcctgcacacgtacgagactctgagaagcaagcacta<br>caatattgcgtttgagctgatcggcctgatttttcttgcctggtacgcaccgcctctggtcacctttgct<br>acctatttctgctttatccatagccgtcgtcatttctcttttgtttggaaacagcttcaacacatgtcct<br>ctaagaaaatgatgataggctccgcgattatcctgagctgcacctcctggttgatcggcggtggtattta<br>ctttttcctgaacagcaaaatgattgcgagcgaagctgcgctgcaaaccgtgtttattggcctggctgcg<br>ctgaccgttccgcacatgatcttgatcgacttcattttccgtccgcactccagccgcattaagatcaaga<br>act**aagctt** |
| pSCAp1.1e vector (pMF821) | gaattcagctgttgacaattaatcatcggctcgtataatgtgtggaattgtgagcggataacaatttcac<br>acaggaaacactcgagtacgtctcaatctctatagagaccttacagctagctcagtcctaggtattatgc<br>tagcgctatgaccatggtacaaagaggagaaaggacatgggtgacccgaaactgctgctggaagttccgc<br>acgctatctcttcttctccgggtaactctctggctgttgttctgcaacgtcgtgactgggaaaacccggg<br>tgttacccagctgaaccgtctggctgctcacccgccgttcgcttcttggcgtaactctgaagaagctcgt<br>accgaccgtccgtcccagcagctgcgttctctgatccgtctgctgaccaaaccggaacgtaaactgtctt<br>ggctgctgccgccgctgtctaacaactaacgcaaaaaaggtctcatgttttagagagacgcaagcttgcc<br>aggcatcaaataaaacgaaaggctcagtcgaaagactgggcctttcgttttatctgttgtttgtcggtga<br>acgctctcctgagtaggacaaatccgcagacgaaagggcctcgtgatacgcctatttttataggttaatg<br>tcatgataataatggtttcttagacgtcaggtggcacttttcggggaaatgtgcgcggaacccctatttg<br>tttatttttctaaatacattcaaatatgtatccgctcatgagacaataaccctgataaatgcttcaataa<br>tattgaaaaaggaagagtatgagtattcaacatttccgtgtcgcccttattcccttttttgcggcatttt<br>gccttcctgtttttgctcacccagaaacgctggtgaaagtaaaagatgctgaagatcagttgggtgcacg<br>agtgggttacatcgaactggatctcaacagcggtaagatccttgagagttttcgccccgaagaacgtttt<br>ccaatgatgagcacttttaaagttctgctatgtggcgcggtattatcccgtattgacgccgggcaagagc<br>aactcggtcgccgcatacactattctcagaatgacttggttgagtactcaccagtcacagaaaagcatct<br>tacggatggcatgacagtaagagaattatgcagtgctgccataaccatgagtgataacactgcggccaac<br>ttacttctgacaacgatcggaggaccgaaggagctaaccgcttttttgcacaacatgggggatcatgtaa<br>ctcgccttgatcgttgggaaccggagctgaatgaagccataccaaacgacgagcgtgacaccacgatgcc<br>tgtagcaatggcaacaacgttgcgcaaactattaactggcgaactacttactctagcttcccggcaacaa<br>ttaatagactggatggaggcggataaagttgcaggaccacttctgcgctcggcccttccggctggctggt<br>ttattgctgataaatctggagccggtgagcgtgggtcacgcggtatcattgcagcactggggccagatgg<br>taagccctcccgtatcgtagttatctacacgacggggagtcaggcaactatggatgaacgaaatagacag<br>atcgctgagataggtgcctcactgattaagcattggtaactgtcagaccaagtttactcatatatacttt<br>agattgatttaaaacttcatttttaatttaaaaggatctaggtgaagatcctttttgataatctcatgac<br>caaaatcccttaacgtgagttttcgttccactgagcgtcagaccagtaagacgggtaagcctgttgatga<br>taccgctgccttactgggtgcattagccagtctgaatgacctgtcacgggataatccgaagtggtcagac<br>tggaaaatcagagggcaggaactgctgaacagcaaaaagtcagatagcaccacatagcagacccgccata<br>aaacgccctgagaagcccgtgacgggcttttcttgtattatgggtagtttccttgcatgaatccataaaa<br>ggcgcctgtagtgccatttacccccattcactgccagagccgtgagcgcagcgaactgaatgtcacgaaa<br>aagacagcgactcaggtgcctgatggtcggagacaaaaggaatattcagcgatttgcccgagcttgcgag<br>ggtgctacttaagcctttagggttttaaggtctgttttgtagaggagcaaacagcgtttgcgacatcctt<br>ttgtaatactgcggaactgactaaagtagtgagttatacacagggctgggatctattctttttatctttt<br>tttattctttctttattctataaattataaccacttgaatataaacaaaaaaaacacacaaaggtctagc<br>ggaatttacagagggtctagcagaatttacaagttttccagcaaaggtctagcagaatttacagataccc<br>acaactcaaaggaaaaggactagtaattatcattgactagcccatctcaattggtatagtgattaaaatc<br>acctagaccaattgagatgtatgtctgaattagttgttttcaaagcaaatgaactagcgattagtcgcta<br>tgacttaacggagcatgaaaccaagctaattttatgctgtgtggcactgctcaaccccacgattgaaaac<br>cctacaaggaaagaacggacggtatcgttcacttataaccaatacgctcagatgatgaacatcagtaggg<br>aaaatgcttatggtgtattagctaaagcaaccagagagctgatgacgagaactgtggaaatcaggaatcc<br>tttggttaaaggctttgagattttccagtggacaaactatgccaagttctcaagcgaaaaattagaatta<br>gtttttagtgaagagatattgccttatcttttccagttaaaaaaattcataaaatataatctggaacatg<br>ttaagtcttttgaaaacaaatactctatgaggatttatgagtggttattaaaagaactaacacaaaagaa<br>aactcacaaggcaaatatagagattagccttgatgaatttaagttcatgttaatgcttgaaaataactac<br>catgagtttaaaaggcttaaccaatgggttttgaaaccaataagtaaagatttaaacacttacagcaata<br>tgaaattggtggttgataagcgaggccgcccgactgatacgttgattttccaagttgaactagatagaca<br>aatggatctcgtaaccgaacttgagaacaaccagataaaaatgaatggtgacaaaataccaacaaccatt<br>acatcagattcctacctacataacggactaagaaaaacactacacgatgctttaactgcaaaaattcagc<br>tcaccagttttgaggcaaaatttttgagtgacatgcaaagtaagtatgatctcaatggttcgttctcatg<br>gctcacgcaaaaacaacgaaccacactagagaacatactggctaaatacggaaggatctgaggttcttat<br>ggctcttgtatctatcagtgaagcatcaagactaacaaacaaaagtagaacaactgttcaccgttacata<br>tcaaaggtgcctaatgagtgaactcacattaattgcgttgcgctcactgcccgctttccagtcgggaaac<br>ctgtcgtgccagctgcattaatgaatcggccaacgcgcggggagaggcggtttgcgtattgggcgccagg<br>gtggtttttcttttcaccagtgagacgggcaacagctgattgcccttcaccgcctggccctgagagagtt<br>gcagcaagcggtccacgctggtttgccccagcaggcgaaaatcctgtttgatggtggttaacggcgggat<br>ataacatgagctgtcttcggtatcgtcgtatcccactaccgagatatccgcaccaacgcgcagcccggac<br>tcggtaatggcgcgcattgcgcccagcgccatctgatcgttggcaaccagcatcgcagtgggaacgatgc<br>cctcattcagcatttgcatggtttgttgaaaaccggacatggcactccagtcgccttcccgttccgctat<br>cggctgaatttgattgcgagtgagatatttatgccagccagccagacgcagacgcgccgagacagaactt |

| | |
|---|---|
| | aatgggcccgctaacagcgcgatttgctggtgacccaatgcgaccagatgctccacgcccagtcgcgtac<br>cgtcttcatgggagaaaataatactgttgatgggtgtctggtcagagacatcaagaaataacgccggaac<br>attagtgcaggcagcttccacagcaatggcatcctggtcatccagcggatagttaatgatcagcccactg<br>acgcgttgcgcgagaagattgtgcaccgccgctttacaggcttcgacgccgcttcgttctaccatcgaca<br>ccaccacgctggcacccagttgatcggcgcgagatttaatcgccgcgacaatttgcgacggcgcgtgcag<br>ggccagactggaggtggcaacgccaatcagcaacgactgtttgcccgccagttgttgtgccacgcggttg<br>ggaatgtaattcagctccgccatcgccgcttccactttttcccgcgttttcgcagaaacgtggctggcct<br>ggttcaccacgcgggaaacggtctgataagagacaccggcatactctgcgacatcgtataacgttactgg<br>tttcacattcaccaccctgaattgactctcttccgggcgctatcatgccataccgcgaaaggttttgcac<br>cattcgatgg |
| PLtetO-1/TetR fragment<br><br>**PvuI/XhoI in bold** | aacctgtcgtgc**cagctg**TTAAGACCCACTTTCACATTTAAGTTGTTTTTCTAATCCGCATATGATCAAT<br>TCAAGGCCGAATAAGAAGGCTGGCTCTGCACCTTGGTGTTCAAATAATTCGATAGCTTGTCGTAATAATG<br>CCGGCATACTATCAGTAGTAGGTGTTTCCCTTTCTTCTTTAGCGACTTGATGCTCTTGATCTTCCAATAC<br>GCAACCTAAAGTAAAATGCCCCACAGCGCTGAGTGCATATAATGCATTCTCTAGTGAAAAACCTTGTTGG<br>CATAAAAAGGCTAATTGATTTTCGAGAGTTTCATACTGTTTTTCTGTAGGCCGTGTACCTAAATGTACTT<br>TTGCTCCATCGCGATGACTTAGTAAAGCACATCTAAAACTTTTAGCGTTATTACGTAAAAAATCTTGCCA<br>GCTTTCCCCTTCTAAAGGGCAAAAGTGAGTATGGTGCCTATCTAACATCTCAATGGCTAAGGCGTCGAGC<br>AAAGCCCGCTTATTTTTTACATGCCAATACAATGTAGGCTGCTCTACACCAAGCTTCTGGGCGAGTTTAC<br>GGGTTTTTAAACCTTCGATTCCGACCTCATTAAGCAGCTCTAATGCGCTGTTAATCACTTTACTTTTATC<br>TAATCTAGACATgttgacggttctccaaaacaaaaagggtacctagccactatagaccaagctgttcatt<br>tttgctttaaagccccgaaattcgattcaccttcccagctttttagttttcaatgtacgcgtccctatca<br>gtgatagagattgacatccctatcagtgatagagatactgagcacatcagcaggacgcactgacc**ctcga**<br>**g**atataatcgaaccatatggga |
| oMFseq45 | GTTTCACATTCACCACCCTGAATTGACTC |
| oMFseq47 | CTTTCGACTGAGCCTTTCGTTTTATTTGAT |
| crtYcYd-F | CAATTTCACACAGGAAACACTCGAGGAGGATTACATatgacttatgctaggttcct |
| crtYcYd-R | gttttatttgatgcctggcaagcttaggacgcgcgagctt |
| oMF361 | ccttggtacaatgctagcctcacacaggaaacactcgaggag |
| oMF362 | cgccatataattgctgtaccGTCGACttaggacgcgcgagctt |
| Strong RBS for Blh (TIR15549) | AGGAGGATTACAA |
| Medium RBS for Blh (TIR1878) | AGCGAGGGGGTTTCCAT |
| Weak RBS for Blh (TIR135) | AGATATAATCGAACCAT |

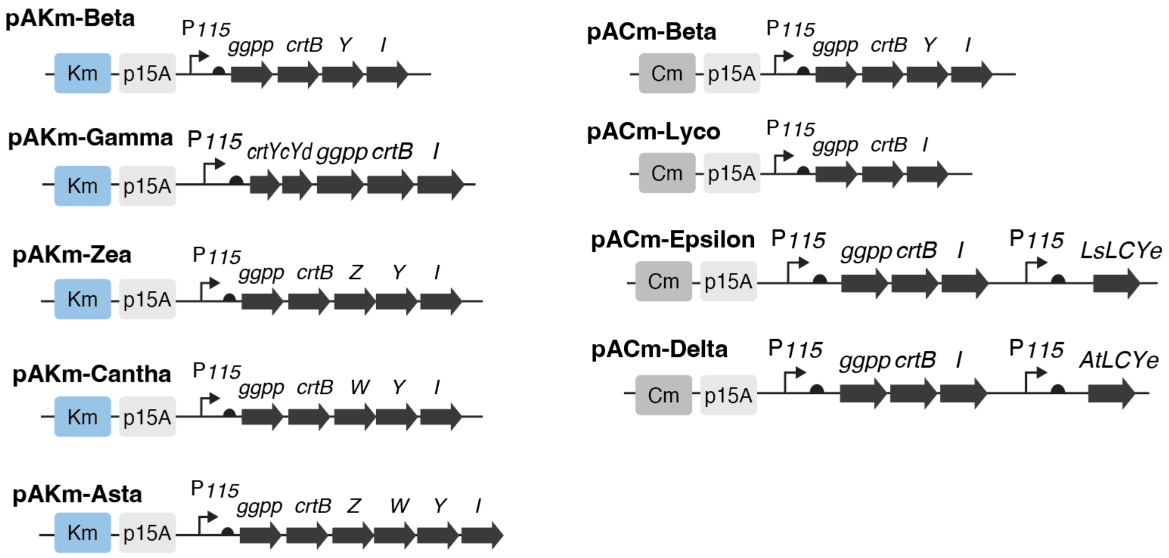


**Supplementary Fig. 1.** Schematics of the plasmid constructs used for carotenoid production. All plasmids except for pAKm-Gamma are derived from Ref. 38 in the main text. See Supplementary Table 1 for details.

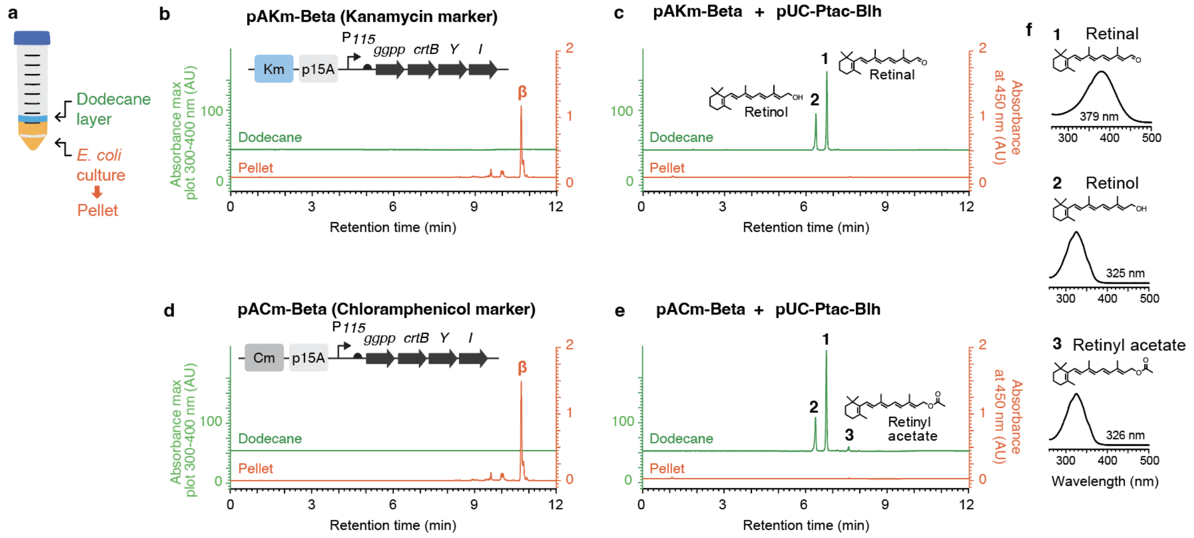


**Supplementary Fig. 2.** Blh cleavage activity of β-carotene in *E. coli* using either chloramphenicol or kanamycin marker. (**A**) Schematic of the *E. coli* culture condition. Dodecane overlay was added on top of *E. coli* culture. The dodecane layer was collected and characterized by UPLC (green chart). The *E. coli* cultures were harvested, and the pellet was subjected to acetone extraction and characterized by UPLC (orange chart). (**B, C**) Carotenoid and retinoid production using kanamycin vector pAKm-Beta. (**D, E**) Carotenoid and retinoid production using chloramphenicol vector pACm-Beta. (**F**) Absorbance spectrum of peaks indicated in panels **C** and **E**.
